# Oncogenic signalling is coupled to colorectal cancer cell differentiation state

**DOI:** 10.1101/2022.04.07.487491

**Authors:** Thomas Sell, Christian Klotz, Matthias M. Fischer, Rosario Astaburuaga-García, Susanne Krug, Jarno Drost, Hans Clevers, Markus Morkel, Nils Blüthgen

## Abstract

Colorectal cancer progression is intrinsically linked to stepwise deregulation of the intestinal differentiation trajectory. In this process, sequential mutations of APC/Wnt, KRAS, TP53 and SMAD4 stepwisely enable an oncogenic signalling network. Here, we developed a novel mass cytometry antibody panel to analyse colorectal cancer cell differentiation and signalling in human isogenic colorectal cancer organoids and in patient-derived cultures. We define a differentiation axis following EphrinB2 abundance in all tumour progression states from normal to cancer. We show that during colorectal cancer progression, oncogenes decrease dependence on external factors and shape distribution of cells along the differentiation axis. In this regard, subsequent mutations can have stem cell-promoting or restricting effects. Individual nodes of the signalling network remain coupled to the differentiation state, regardless of the presence of oncogenic signals. Our work underscores the key role of cell plasticity as a hallmark of cancer that is gradually unlocked during colorectal cancer progression.

## Introduction

Colorectal cancer (CRC) is one of the most prevalent neoplastic diseases. It progresses by cumulative acquisition of mutations in a number of specific oncogenes and tumor suppressors, each modulating one or more intracellular signalling cascades.^1^ Activation of the Wnt/β-Catenin pathway through functional loss of APC is the most common initiating event for most colorectal adenomas, and plays a pivotal role in promoting a crypt progenitor-like phenotype.^2,3^ Constitutive activation of the EGFR/MAPK pathway are commonly found in adenomas and carcinomas, and activating mutations of the pathway members KRAS and its downstream kinase BRAF are found in 40 % and 5–10 % of CRCs, respectively.^4,5^ Another common alteration is functional loss of TP53, disrupting DNA damage response and repair. As this mutation is commonly found in colorectal carcinoma but not in adenoma, it is closely tied to the adenomacarcinoma transition.^3,6,7^ Impaired transforming growth factor β (TGF-β) pathway function is a rather late event in cancer progression, and the mediator SMAD4 is inactivated in 10–15 % of CRCs.^3^

While the mutational progression of CRC from adenomas to carcinomas is well studied, it is still unknown how the key events shape cell-type specific signal transduction and differentiation states. Recent advances in single cell technologies and organoid model systems now allow studying cell-state specific effects of oncogenes.^8,9^ For example, we have recently combined transgenic intestinal organoids and single cell methods to show that KRAS-mediated activation of ERK signalling is restricted to undifferentiated cells.^10^.

In this study, we aimed to investigate how oncogenes and growth factors drive cell signalling, cell plasticity, and differentiation. We employed human colon organoids from healthy donors that were genetically engineered to model canonical adenoma-carcinoma progression.^11^ These organoids preserve the intestinal epithelium’s continuous differentiation through multiple phenotypic states and are thus particularly well suited for unveiling effects of oncogenes on phenotypic plasticity.^12^ We analysed these organoids using mass cytometry (MC), a technique that allows to measure tens of markers in single cells. Analysis of the high-dimensional data set showed specific effects of cell-intrinsic mutations on the signalling network which in turn anchored cells on different positions on a differentiation axis. We verified our findings using tumor-derived organoids.

## Methods

### Antibody panel

We designed an antibody panel targeting a selection of cell-type specific epitopes, multiple cell-state markers, and various members of intracellular signalling cascades (Table 1). Most of our antibodies were conjugated in-house using Maxpar X8 labelling kits, following the manufacturer’s protocol (Fluidigm, 201300).

**Table 1:** Antibody marker panel. All antibodies not purchased from Fluidigm were conjugated in-house. Channels were chosen to minimise potential spillover of high-abundance markers into low abundance channels.

| type | target | vendor | clone | label |
| --- | --- | --- | --- | --- |
| surface | CD133 (PROM1) | Miltenyi | AC133 | 154Sm |
| surface | CD24 | Fluidigm | ML5 | 169Tm |
| surface | CD326 (EpCAM) | BioLegend | 9C4 | 167Er |
| surface | CD44 | Fluidigm | BJ18 | 166Er |
| surface | EphB2 | BD | 2H9 | 158Gd |
| surface | LGR5 | Fluidigm | 4D11F8 | 161Dy |
| surface | PTK7 | Miltenyi | 188B.12.45 | 146Nd |
| intracellular | cCasp3 | Fluidigm | D3E9 | 142Nd |
| intracellular | cPARP | Fluidigm | F21-852 | 143Nd |
| intracellular | IκBα | Fluidigm | L35A5 | 164Dy |
| intracellular | Ki-67 | Fluidigm | B56 | 162Dy |
| intracellular | Krt20 | CST | D9Z1Z | 176Yb |
| intracellular | p-p38 [T180/Y182] | Fluidigm | D3F9 | 156Gd |
| intracellular | p-p53 [S15] | CST | 16G8 | 172Yb |
| intracellular | p4e-BP1 [T37/46] | CST | 236B4 | 170Er |
| intracellular | pAkt [S473] | Fluidigm | D9E | 152Sm |
| intracellular | pChk1 [S345] | CST | 133D3 | 148Nd |
| intracellular | pChk2 [T68] | CST | C13C1 | 141Pr |
| intracellular | pERK1/2 [T202/Y204] | Fluidigm | D13.14.4E | 171Yb |
| intracellular | pH2A.X [S139] | Fluidigm | JBW301 | 147Sm |
| intracellular | pMEK1/2 [S217/221] | CST | 41G9 | 151Eu |
| intracellular | pNF-kB p65 [S536] | CST | 93H1 | 155Gd |
| intracellular | pS6 [S235/236] | Fluidigm | N7-548 | 175Lu |
| intracellular | pSmad1 [S463/S465] / pSmad8 [S465/S467] | BD | N6-1233 | 149Sm |
| intracellular | pSmad2 [S465/467] / pSmad3 [S423/425] | CST | D27F4 | 153Eu |
| intracellular | Vimentin | CST | D21H3 | 165Ho |
| intracellular | YAP | CST | D8H1X | 150Nd |

### Organoid and cell line culture

The following organoid lines were used in this study: Normal colon organoids, established from human intraoperative material (Charité ethics committee approval #EA4/015/13), isogenic normal and CRISPR-modified human colon organoids (Drost et al. 2015^11^), human colon organoid lines OT227, OT302 (Schütte et al. 2017^13^) and P009T, P013T (Uhlitz et al. 2021^14^).

Organoids were cultured in growth-factor reduced Matrigel (Corning, 356230), according to previously published protocols.^8,15^ Culture medium was Advanced DMEM/F12 (Gibco #12634010) supplemented with 1x GlutaMAX (Gibco #35050061), 1x N-2 (Gibco #17502048), 1x B-27 (Gibco #17504044), 500 mM N-Acetylcysteine, 1 M HEPES, 100 U/ml Penicillin-Streptomycin, and 100 µg/ml Primocin (Invitrogen #ant-pm-1). This base medium was further supplemented with a selection of growth factors, inhibitors, and conditioned media which varied by line (Table 2). A, AK, AKP, AKPS media were designed to maintain selection pressure towards the inherited mutations.^11^ R-Spondin conditioned medium was produced using HEK 293T HA-RSpo-1-FC clone 3B, according to previously published protocols.^16^ Wnt-conditioned medium was produced using L cells WNT3A clone 5.5, according to a previously published protocol.^17^

**Table 2:** Custom culture medium conditions for the different lines measured.

| substance | source | conc | NCO | A | AK | AKP | AKPS | PDO |
| --- | --- | --- | --- | --- | --- | --- | --- | --- |
| Wnt CM | in-house | 50 % | • |  |  |  |  |  |
| Rspo1 CM | in-house | 5 % | • |  |  |  |  |  |
| Noggin | Peprotech #250-38 | 100 ng/ml | • | • | • | • |  |  |
| EGF | Peprotech #315-09 | 50 ng/ml | • | • |  |  |  | • |
| Nutlin3 | Cayman #10004372 | 1 μM | | | | • | • | |
| SB 202190 | Sigma-Aldrich #S7067 | 3 μM | • | • | • | • | • | • |
| A 83-01 | Tocris #2939 | 500 nM | • | • | • | • | • | • |

Prior to measurement, small fully formed organoids were re-plated without de-aggregation and kept in base medium supplemented with combinations of Wnt, R-spondin 1, and EGF for 2 or 4 days. During this time, Wnt was supplied as recombinant mouse Wnt3a (100 ng/ml; Time Bioscience).

### Mass cytometry

Organoids were collected from culture plates and dissociated into single cells using a 1:1 mixture of TrypLE Express (Gibco, 12604013) & Accutase (Gibco, A1110501), supplemented with 100 U/ml Benzonase (Pierce, 88700). Subsequently, the single-cell suspension was kept in 5 µM Cisplatin/PBS solution (Fluidigm, 201064) for 5 minutes at 37 °C to stain dead cells and cell debris. After washing twice with culture medium to remove Cisplatin, cells were kept for 30 minutes at 37 °C in their original medium supernatant, which was preserved prior to harvesting the organoids. Following this resting phase, samples were fixed for 10 minutes at room temperature in a mixture of 1:1.4 PBS/BSA 10 % & “Smart Tube” proteomic stabilizer (NC0627333) and subsequently frozen at −80 °C.

One day prior to measurement, samples were thawed and barcoded using the Cell-ID 20-Plex Pd Barcoding Kit (Fluidigm, 201060) according to manufacturer’s instructions. After subsequent pooling of all samples, cells were incubated with metal-isotope tagged antibodies in a two step protocol. First, antibodies targeting cell-surface proteins (Table 1) were added to the cell pool for 30 minutes at room temperature, followed by 10 min fixation in 2 % Formaldehyde at room temperature and membrane permeabilisation in ice-cold methanol. Subsequently, antibodies targeting intracellular and intranuclear proteins were added to the cell pool for 30 minutes at room temperature. In between each step cells were washed in Maxpar Cell Staining Buffer (Fluidigm, 201068). DNA staining was performed using Cell-ID Intercalator-Ir (Fluidigm, 201192a) with a final concentration of 62.5 nM in PBS for 20 minutes at room temperature. After overnight fixation in 2 % Formaldehyde, stained cells were washed once with Cell Staining Buffer, twice with doubly distilled water, and filtered through a 30 µm cell strainer.

EQ™ Four Element Calibration Beads (Fluidigm, 201078) were added 1:10 to the cell suspension prior to analysis in a Helios mass cytometer (Fluidigm). The instrument’s software was used to normalise measured data for machine-related variance based on the added calibration beads. Further data processing and plotting was performed in R^18^ using various packages of the Tidyverse family^19^. We used the CATALYST package for de-convolution of barcoded samples and spillover compensation^20^. Further pre-processing steps involved filtering out EQ bead events, gating for single-cells via event length & DNA parameters, and excluding dead cells via a Platin-Iridium gate. For further data processing and plotting, antibody marker signals were arsinh transformed.

Biological replicates of the CRC progression series experiment were measured in separate MC runs. Because we did not include anchor samples, we instead computed scaling factors to match between runs the 85th percentile of each antibody signal across all perturbation conditions measured, eliminating technical variation while preserving treatment-related effects. We sampled equal numbers of cells per perturbation condition to ensure that the adjustment was not dominated by any specific perturbation. Run 2 was adjusted using run 1 as reference, run 3 was adjusted using the normalised run 2 as reference.

## Results

### An antibody panel targeting the intestinal epithelium

The intestinal epithelium is a complex tissue consisting of a hierarchy of cell types. Therefore, characterisation of cellular signalling in this tissue needs to take into account its different cell types and states. We employed Mass cytometry (MC), also known as Cytometry by time of flight (CyTOF), a high-dimensional single cell proteomics method, to measure cell type and cell state markers and simultaneously quantify the activity of various intracellular signalling pathways.^21,22^ We established a custom MC antibody panel, suitable for simultaneously assessing cell phenotypes and cell signalling states in human colon organoids (Fig. 1a, Table 1).

**Figure 1:**
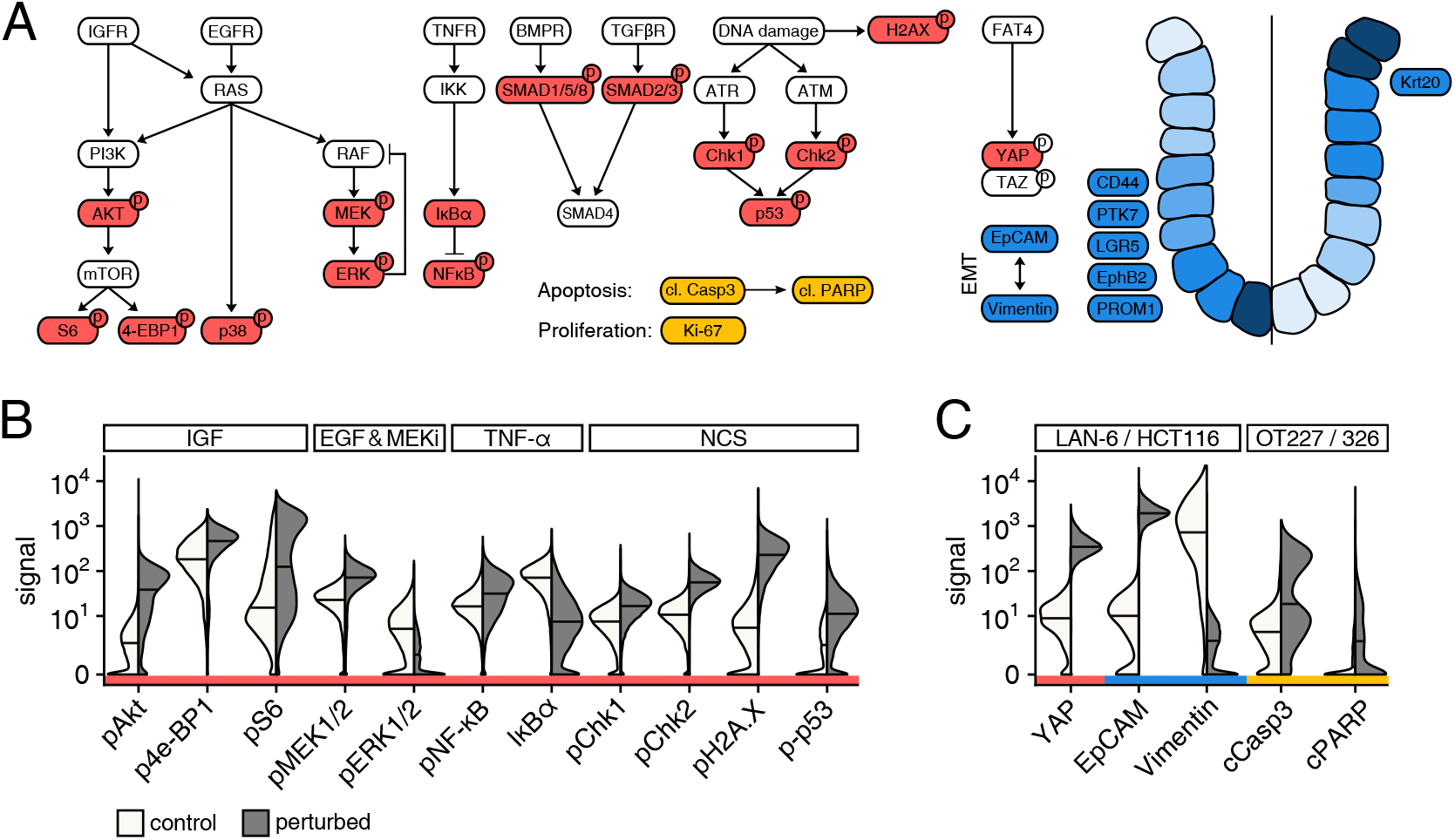
A CyTOF panel to analyse cell type, state, and signalling. **a** Visual representation of the antibody panel.Intracellular signalling markers in red, cell-state markers in blue, cell-type markers in green. **b** Functional control MC experiments for selected antibodies of the signalling sub-panel. Split violin plots with medians. HCT116 cells were subjected to small molecule stimulants & inhibitors to create positive & negative control conditions. Compounds used: IGF (100 ng/ml, 30 min.), EGF (25 ng/ml, 10 min.), AZD6244 (MEK inhibitor; 10 µM, 60 min.), TNF-α (20 ng/ml, 30 min.), Neocarzinostatin (500 ng/ml, 30 min.). **c** Functional control MC experiments for selected antibodies of the cell-state & cell-type specific sub-panels. Split violin plots with medians. Cell lines with previously known differential expression of total proteins in question were used.

Antibodies were functionally tested using small molecule substances on 2D CRC cell lines or by using two different lines with known differential expression of certain protein markers (Fig. 1b,c).^23^ Signals of antibodies targeting phosphorylated Akt, S6, and 4-EBP1 (pAkt, pS6, p4-EPB1) were increased after stimulation of the IGFR/PI3K axis compared to unstimulated controls (Fig. 1b). Phosphorylated MEK (pMEK) was increased after stimulation with EGF and simultaneous inhibition of MEK function, strengthening EGF-induced ERK to MEK signalling by eliminating a well-known negative feedback loop.^24^ Abundance of phosphorylated ERK (pERK) was decreased under these conditions due to upstream MEK inhibition. TNF-α stimulation led to degradation of IκBα and an increased phosphorylation of its inhibition target NF-κB (pNF-κB). DNA damage inflicted by Neocarzinostatin (NCS) increased phosphorylation of both Chk1 and Chk2 as well as other well known DNA damage response markers histone H2AX and p53 (pChk1, pChk2, pH2A.X, p-p53). Hippo pathway factor YAP and epithelial cell marker EpCAM had distinctly higher abundances in CRC line HCT116 compared to mesenchymal neuroblastoma line LAN6 (Fig. 1c). Marker of metastatic progression and epithelial-to-mesenchymal transition, Vimentin was mostly absent in HCT116 cells yet strongly expressed in LAN6. For validation of apoptosis markers cleaved Caspase 3^25,26^ and cleaved PARP^27^, we compared two previously described patient-derived CRC organoid lines OT326 and OT227 that are known to display a large or absent apoptotic population, respectively.^10,13^ Similar to previous experiments, a noticeable number of cells in OT326 but not OT227 had high abundances of both apoptosis markers. Signals of additional cell-type markers LGR5, PROM1, CD24, EphB2, Krt20, and PTK7 all correlated with previously known RNA expression data of the cell lines compared (Fig. S1).^23^ These data establish that our MC antibody panel is well suited for measuring cell phenotypes of intestinal organoids while simultaneously characterising intracellular signalling states.

### EphB2 gradient parallels differentiation axis in normal colon organoids

To assess cell state heterogeneity in non-cancerous conditions, we profiled human normal colon organoids (NCO). We cultured NCOs for 4 days in either complete medium (+Wnt) or Wnt-deprived conditions (−Wnt), subsequently prepared MC samples in two biological replicates, and used sample barcoding to reduce inter-sample variability (Fig. 2a, Suppl. Fig. S2).

**Figure 2:**
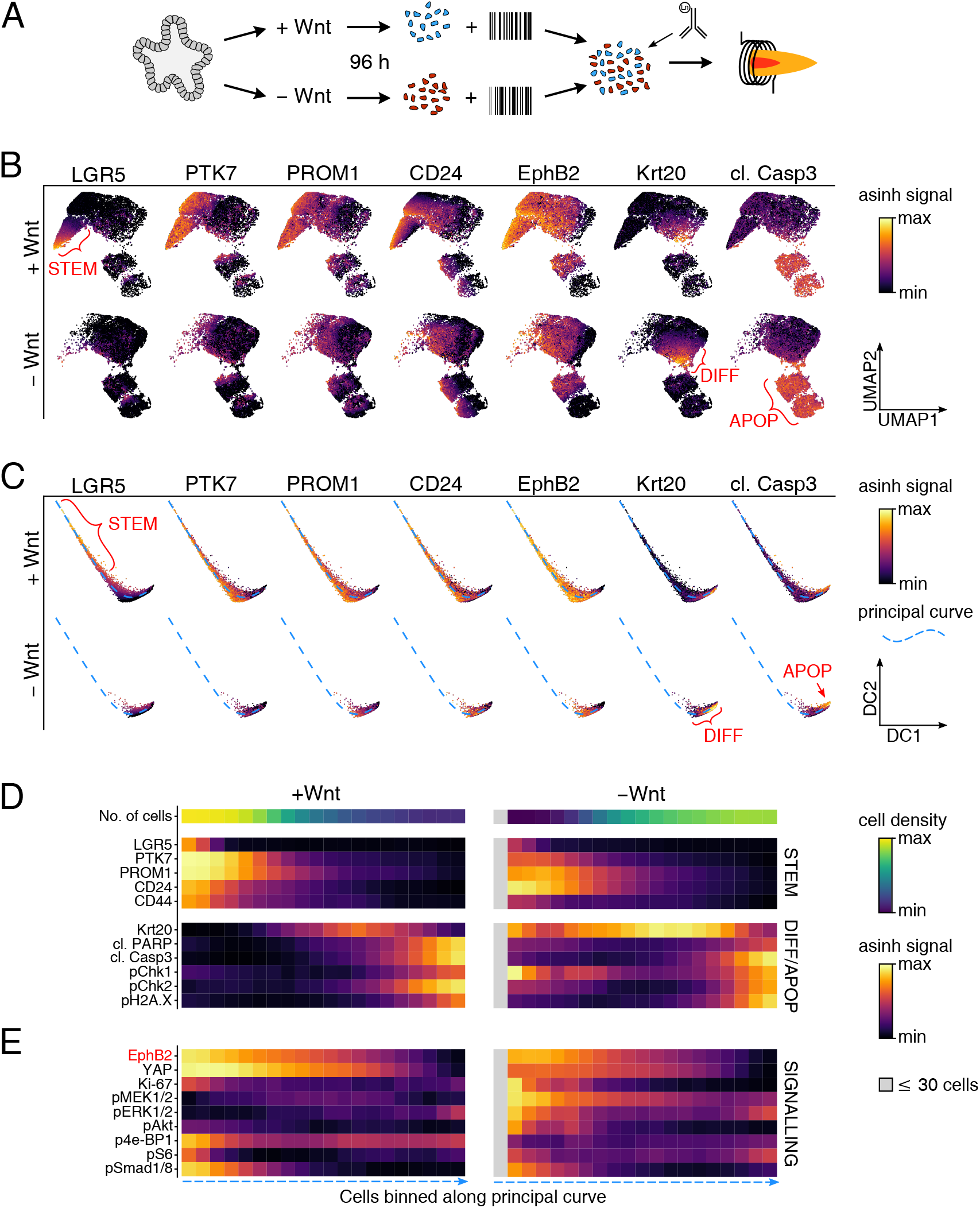
MC reveals differentiation trajectory in normal colon organoids. **a** Experimental workflow. Human normal colon organoids were subjected to ± Wnt conditions for 4 days, subsequently dissociated into single cells, barcoded, pooled, stained with the antibody panel, and measured via mass cytometry. **b** UMAP embedding based on the 7 cell-type markers shown. Colour scale depicts normalised marker signals in ± Wnt conditions. Stem cell, differentiated, apoptotic regions are marked in red. **c** Diffusion map embedding based on the cell-type markers also used in (b). Fitted principal curve of the diffusion map space is depicted as a dashed blue line. **d/e** Heatmap showing cell distribution and mean protein marker signals within equally sized bins along the principal curve defined in (c). Means rescaled per marker and smoothed out using a 3-bin wide running average. Bins containing less than 30 cells were excluded and are shown in grey.

Dimension reduction using uniform manifold approximation and projection (UMAP) of seven cell-type specific markers yielded a meaningful clustered representation of cell phenotypes^28^ (Fig. 2b). High signal intensities of LGR5 and PTK7, two markers associated with the intestinal stem-cell niche,^29,30^ marked a distinct area from regions that were high in Krt20, a marker expressed in differentiated villus regions.^31^ EphB2, known to be closely linked to the crypt-villus axis,^32^ gradually decreased between the two extremes. Apoptotic cells, marked by high abundance of cleaved Caspase 3, formed two separate clusters near the region of differentiated cells. Under conditions of Wnt withdrawal, we observed a decrease of cells within the stem/crypt region and an increase in the area representing differentiated or apoptotic cells. This is expected as an external Wnt source is required for normal colon stem cell maintenance^2^. We concluded that the cell-type markers used to create this UMAP contained sufficient information to cluster the most relevant cell differentiation stages of colon organoid cells.

To define a continuous differentiation trajectory within the dataset, we re-embedded cell events into a diffusion map, using the same seven cell-type markers. Unlike UMAPs which emphasise clustering of high-dimensional data, diffusion maps preserve transitions between neighbouring points.^33^ In the space defined by diffusion components 1 and 2, cells aligned on an almost one dimensional path (Fig. 2c). The phenotypical regions of cells already described in Fig. 2b were also visible in this representation, with stem cells clustering at one end of the path and a differentiated region at the other end, marked with high Krt20 levels. Caspase 3, marking apoptotic cells, was enriched at the very end of the trajectory. Similar to the UMAP embedding, cells of the −Wnt conditions were shifted towards the differentiated region. As this diffusion map faithfully reproduces the intestinal epithelium’s inherent differentiation trajectory, we fitted a principal curve (blue dashed line) that defines a pseudo-time axis on which intestinal organoid cells move along towards differentiation.

We sorted and divided the dataset into bins along the previously defined pseudo-time differentiation axis to assess the distribution of cells along this trajectory as well as the dynamics of protein markers (Fig. 2d). In full-medium (+Wnt) conditions, most cells were in the left-sided bins, representing early, stem-like phenotypes. In contrast, in −Wnt conditions cells shifted towards the right-sided bins that contained more differentiated cells. Interestingly, protein marker distribution along the axis was rather constant across conditions. Under both conditions, markers typically associated with stem-like, proliferating cells such as LGR5, PTK7, and PROM1 were tied to the undifferentiated left side of the pseudo-differentiation axis. Krt20 was most prominently expressed at the two-thirds point whereas markers of apoptosis and DNA damage response, such as cleaved Caspase 3, pChk1/2, and pH2A.X, had their peak at the very end of the binning axis. Unexpectedly, intracellular signalling markers were largely linked to differentiation state as described by the principal curve axis (Fig. 2e). Ki-67, p4e-BP1, and pS6 were mostly active in undifferentiated cells, indicating cell proliferation and high metabolic activity in this region. The Hippo pathway downstream transcription factor YAP was restricted to the beginning of the pseudo-differentiation axis. In contrast, activity of the MEK/ERK pathway was partly uncoupled from cell differentiation, as Wnt withdrawal led to increased phosphorylation of both markers in undifferentiated cells.

Taken together, the mass cytometry analyses of normal colon organoids shows that cell signalling and cell phenotypes are strongly linked in normal colon organoids. Notably, EphB2 correlated especially well with the pseudo-time differentiation axis (Fig. 2e), and we therefore concluded that EphB2 is a suitable marker for the pseudo-differentiation axes, as noted before.^32^

### Oncogenes modulate cell phenotype to facilitate CRC progression

To assess interplay between cell phenotypes, signalling, and oncogenic mutations in CRC progression, we employed a previously established set of four isogenic human colon organoid lines derived from NCO using sequential genetic modifications.^11^ These represent the canonical step-wise progression of CRC, namely APC loss, KRAS activation, TP53 loss, and SMAD4 loss and thus termed, NCO-A, NCO-AK, NCO-AKP, NCO-AKPS, respectively. We perturbed ligand-based signalling by growing all organoids in full medium containing Wnt, or medium lacking Wnt and/or EGF for 48 h and subsequently performed MC analysis (Fig. 3a).

**Figure 3:**
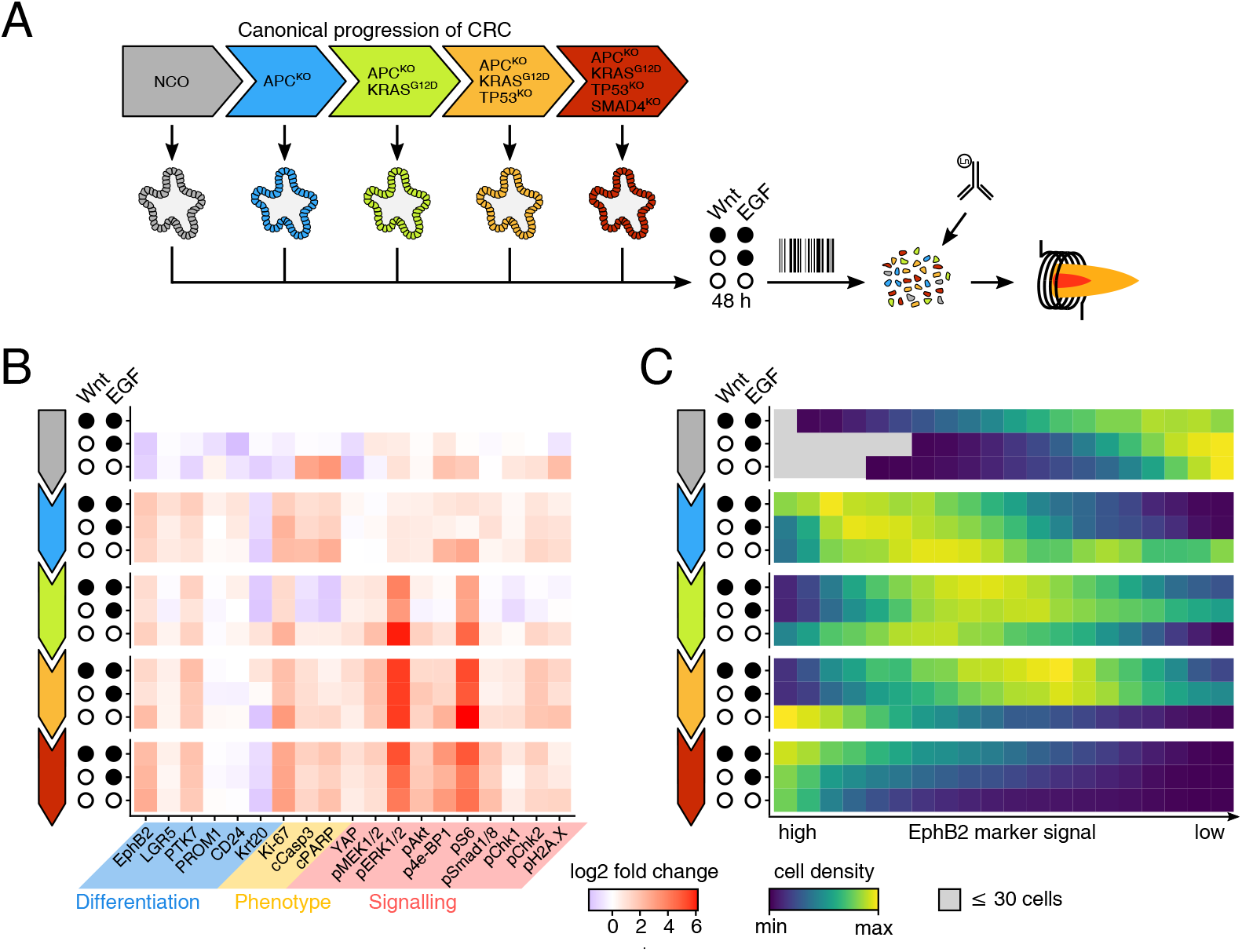
CRC progression model shows non-continuous path towards stemness. **a** Experimental workflow. Human normal colon organoids, sequentially mutated with key oncogenes of canonical CRC progression (APC LoF, KRAS GoF, p53 LoF, SMAD4 LoF)^11^ were subjected to three different media conditions (complete, −Wnt, −Wnt/−EGF) and processed into one multiplexed MC sample. **b** Log2 fold changes per marker relative to complete medium NCO condition. Based on multiplexed mass cytometry sample, showing all 5 sequentially mutated lines described in (a). **c** Cell density distributions along a pseudo-differentiation axis defined by Ephrin B2 marker signal. Normalised by medium & line. High Ephrin B2 (stem/TA region) on the left and low Ephrin B2 (differentiated/apoptotic region) on the right.

As before, the NCO line reacted to Wnt withdrawal with a decrease of stem cell and crypt markers such as LGR5, PTK7, and EphB2 (Fig. 3b and S4). Additional EGF starvation resulted in a decrease of Krt20 while DNA damage and apoptosis markers cleaved Casp3, pChk1/2, and pH2A.X were increased. In NCO-A, APC loss led to increased abundances of stem cell and crypt markers while simultaneously reducing Krt20, in line with the well-established role of Wnt/β-Catenin signals in stabilising crypt progenitor fate.^2^ Increased Ki-67 levels also indicated higher proliferation in the absence of functional APC. Wnt withdrawal had no effect on these or derived organoids, compatible with cell-autonomous activation of Wnt/β-Catenin signalling after APC loss. NCO-AK, containing oncogenic KRAS in addition to APC loss, showed increased abundance of pMEK, pERK, and pS6, but notably not of pAkt and p4e-BP1. Levels of intestinal stem cell marker LGR5 decreased and returned to levels of the unmutated NCO. Paradoxically, EGF starvation led to increased pERK in NCO-AK, as well as increased levels of Krt20, cleaved Casp3, and DNA damage markers. NCO-AKP, with an additional TP53 mutation, showed variable effects on pChk1, pChk2, and pH2A.X across replicates, suggesting a further dependence on external culture conditions (Fig. S4). Finally, SMAD4 knock-out in NCO-AKPS showed increased abundances of related co-factors SMAD1 and SMAD8. Taken together, all CRC driver mutations showed direct effects on downstream signal transduction, but also caused more global changes in signalling and cell type markers.

Based on the relationship between EphB2 and differentiation state of cells established above (Fig. 2d), we divided this dataset into 20 equally sized bins after sorting by EphB2 abundance (Fig. 3c). When comparing cell lines, we found that mutations modified cell distributions along the pseudo-differentiation axis. NCO predominantly occupied the bins on the right side, corresponding to more differentiated cells, while NCO-A inhabited the left side of the axis. Subsequent KRAS mutation in NCO-AK partly reversed this effect towards more differentiation, and cells of NCO-AKP maintained this position. SMAD4 loss had the most drastic effect, causing an almost uniform stem/progenitor-like phenotype with high expression of EphB2. Removal of EGF from the culture medium also affected cell phenotypes, promoting differentiation prior to KRAS mutation and stemness thereafter. Overall this analysis shows that different oncogenic mutations and external growth factors both shape cell differentiation states and that growth factors can have opposing effects contingent on the presence of oncogenes.

### Differentiation state and oncogenes account for most variance in CRC

Within the dataset based on oncogene and extrinsic signal perturbations (Fig. 3), cells can be categorised using three dimensions: Mutational state, available growth factors, and differentiation phenotype (Fig. 4a). To disentangle how intracellular signalling states are informed by these dimensions, we modelled binned marker abundances using analyses of variance (ANOVA). The ANOVA model contained mutational state, growth factors, and the EphB2 axis position as factors. We also included an interaction term between mutational state and growth factors to account for the divergent effects of growth factors depending on oncogenes, as seen above. We trained the model on three biological replicates of this experiment (Fig. 4b, for replicates see Fig. S4).

**Figure 4:**
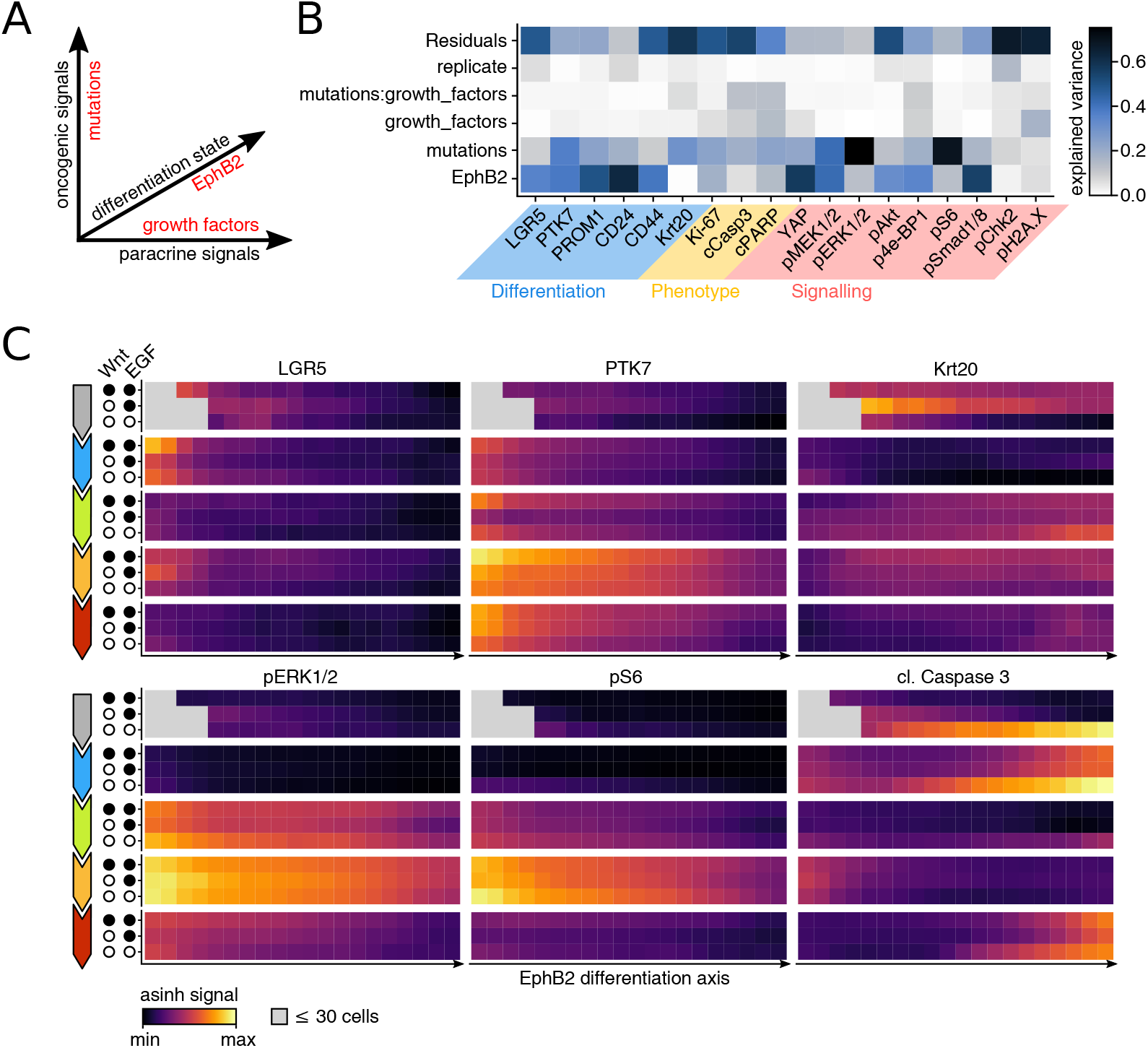
Many signalling nodes are linked to CRC organoid cell differentiation states. **a** Dimensions of the dataset shown. Cells are mostly defined by oncogenes (intracellular signals), growth factors in the culture medium (paracine signals), and position on the EphB2 pseudo-differentiation axis. **b** Analysis of variance per marker, using the dimensions described in (a), an interaction term of mutations and supplied growth factors, as well as the replicate ID as linear model factors. Colour scale shows fraction of variance explained per predictor. **c** Mass cytometry signals of selected marker proteins along a pseudo-differentiation axis defined by Ephrin B2 marker signal. Organoid lines as row blocks and medium conditions as intra-block rows. Colour scale depicts normalised mean marker signal across three replicates.

Across most markers, variance explained by replicates was generally very low, indicating a high reproducibility (Fig. 4b). Markers of stem cell or crypt progenitor phenotypes LGR5, PTK7, PROM1, CD24, CD44, YAP were all strongly linked to the EphB2 pseudo-differentiation axis, and modulated by mutations. In contrast, markers of MAPK signalling and proliferation (pERK1/2, pMEK1/2, pS6, and Ki-67) were strongly explained by the mutational state. Only few markers were influenced by the presence of growth factors, most notably cleaved Caspase 3 which mainly indicated apoptosis in non-KRAS mutated −EGF conditions. This was reflected by the highest percentage of variance explained by the interaction term between mutational state and growth factors across all markers.

These patterns could also be visualised by mean marker abundance per EphB2 bin (Fig. 4c, Fig. S5). For example, LGR5 and PTK7 having graded signals along the EphB2 axis, while also varying greatly between cell lines. LGR5 was mainly present in EphB2 high cells of the APC mutant and APC/KRAS/TP53 mutant lines, whereas PTK7 was stronger in lines carrying a TP53 mutation. Variance of Krt20 was mainly explained by APC mutation. MAPK signal transducers, in contrast, showed a strong increase after KRAS activation and further after subsequent TP53 loss, while SMAD4 knockout diminished proliferation marker signals. Overall, both the analyses of variance and inspection of single marker heatmaps show that most signalling states in CRC cells are determined by mutation and differentiation state, and are largely decoupled from external signals like EGF and Wnt.

### Tumours are also mainly defined by differentiation state and oncogenes

Unlike our isogenic CRC progression model lines, which inherit a defined mutational profile, genotypes of individual patient-derived tumour organoids are complex and heterogeneous. To investigate if our finding that signalling is largely determined by oncogenic mutations and cell states extends to human CRC, we employed four well characterised patient-derived organoid lines (OT227, OT302, P009T, and P013T; Fig. 5a).^13,14^ We therefore cultured the lines in the presence or absence of Wnt and/or EGF (Fig. 3a), and observed similar effects on the distribution of cells across a pseudo-differentiation axis defined by EphB2 (Fig. 5b). Both OT227 and P013T cells underwent differentiation in response to Wnt withdrawal, but de-differentiated when both Wnt and EGF were removed. In contrast, OT302 and P009T did not show growth factor dependencies.

**Figure 5:**
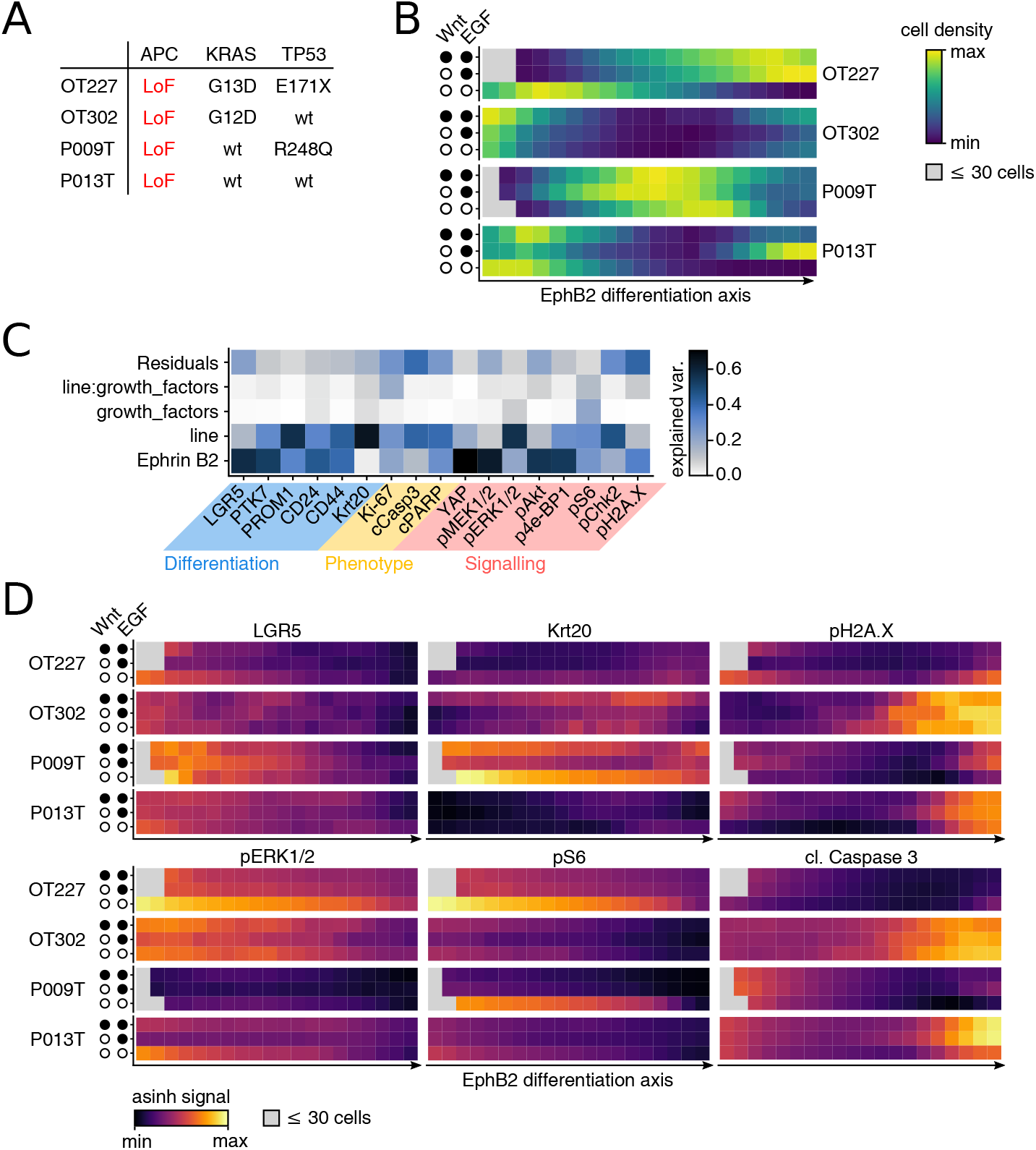
Patient-derived CRC organoid signalling networks are largely defined by differentiation state. **a** Mutational state of oncogenes APC, KRAS, and TP53 for patient-derived organoid lines shown in this figure. **b** Cell distributions per line & medium along a pseudo-differentiation axis defined by Ephrin B2 marker signal across all 4 lines. **c** Analysis of variance per marker, using the dimensions described in Fig. 4a as well as an interaction term of mutations and supplied growth factors as linear model factors. Colour scale shows fraction of variance explained per predictor. **d** Mass cytometry signals of selected marker proteins along a pseudo-differentiation axis defined by Ephrin B2 marker signal. Organoid lines as row blocks and medium conditions as intra-block rows.

We again performed analyses of variance for single markers to identify dynamic signals within the dimensions defined in Fig. 4a. As previously, cell-type markers but also many signalling markers were strongly dependent on the position along the pseudo-differentiation axis defined by EphB2. However, line-dependent differences were generally much larger in this dataset, while growth factors affected only a small number of signal transducers, such as pERK and pS6 (Fig. 5c). In line with the ANOVA, variance in stem-cell and crypt-region associated markers LGR5, PTK7, PROM1, CD24, CD44, and YAP was regionalised on the EphB2 pseudodifferentiation axis (Fig. 5d, Fig. S6). Expectedly, LGR5 was associated with early pseudodifferentiation axis position in all lines; however unexpectedly Krt20 was not limited to late positions of the axis. DNA damage response markers such as pH2A.X and pChk2 were restricted to late positions of the TP53 wildtype lines OT302 and P013. Apoptosis as indicated by cleaved Caspase 3 closely mimicked pH2A.X activity in being associated with a differentiated cell state. While ERK phosphorylation was generally higher in KRAS-mutated lines OT227 and OT302, it was further elevated by EGF starvation in OT227 cells, similar as seen above in NCO-AK. Collectively, this data shows that the principle of differentiation state and mutations dictating signalling activity in single CRC cells extends to complex patient-derived organoids.

## Discussion

Human colon organoids are a powerful model system for studying signalling and phenotypes of cells in the context of a complex epithelial tissue in health and disease.^34^ Here, we analysed human organoids with single cell resolution to study how oncogenes and extra-cellular signals affect cell-state distribution, and in turn how cell states affect signal transduction. Mass cytometry (MC) enabled us to investigate intracellular signalling and cell state markers at the same time. We found that EphB2 defines a pseudo-differentiation axis in normal and CRC tissue, allowing us to investigate signalling changes along this axis. Using a series of isogenic cell lines of key canonical adenoma-carcinoma progression steps, we could compare signalling at various stages of colorectal cancer development. We found that the cell differentiation state was the most important determinator of cell signalling state and that oncogenes restricted permissible cell differentiation states, thus limiting the impact of extrinsic Wnt and MAPK effectors.

Wnt signalling is known to be essential for maintaining intestinal stem and crypt cell states, and EphB2 is a known graded marker of crypt cells.^2,32^ Ephrins, including EphB2, are thought to be Wnt targets and repressors of CRC progression.^35^ However, we found that EphB2 expression remained graded also in organoid lines with engineered loss of the Wnt signal regulator APC, suggesting that signalling pathways beyond Wnt play a role in the regulation of EphB2. Interestingly, the Hippo transducer YAP correlated well with EphB2 expression and could therefore be a candidate regulator compatible with its role to regulate stem-cell fate in the colon^36^ (Fig. 2e). High EphB2 abundance remained associated with stem-cell markers and stem-cell specific cell signalling network states throughout pre-cancerous, and cancerous progression states, shedding further light on EphB2’s ability to predict CRC relapse in patient cohorts.^37^

Our analysis shows that oncogenic mutations shift cells along the differentiation axis, which also largely determines signalling state. Each of the successive mutations in APC, KRAS, TP53, and SMAD4 had a profound effect on cell differentiation as defined by the pseudodifferentiation axis. It was somewhat surprising however that cells did not gradually shift towards higher stemness, but were subject to sometimes opposing effects. While the Wnt effector APC moved cells towards high expression of stem-cell markers, mutation of the MAPK signal transducer KRAS promoted differentiation. Antagonistic Wnt and MAPK activity has recently been shown to limit anti-MAPK therapy efficiency in CRC.^14,38^ Our data extend this antagonistic model to activities of the APC and KRAS driver mutations during CRC progression. Additional loss of TGF-βand BMP signal transducer SMAD4 reversed the KRAS effect in our data. Loss of SMAD4 could thus contribute to progression of CRC by promoting cancer cell plasticity towards stemness that was restricted by prior MAPK activating mutations. Interestingly, while consecutive mutations also shift mouse CRC organoids towards stem-cell fate in a recently published MC dataset^9^ (Fig. S7), the effect of KRAS activation on stemness appears to be different between human and mouse organoid models. These potential species differences highlight the importance of such studies in human model systems. Collectively, our results suggest that APC, KRAS, and SMAD4 mutations play different and partially opposing roles in controlling cell plasticity, which was recently recognised as a new hallmark of cancer.^39^

We found that cells inhabiting similar positions along the differentiation axis shared many signalling network states across organoids of the isogenic progression series. Not only stemcell markers but also signal transducers such as p-MEK, p-Akt, p-4ABP1, pSmad1/8, and YAP were strongly correlated to the EphB2 axis (Fig. 4b). Indeed, p-MEK, p-Akt, p-4ABP1, p-S6, and YAP were also strongly correlated to the EphB2 axis in patient-derived lines (Fig. 5c). Therefore, our data suggests that oncogenes and tumour suppressors cannot uncouple parts of the cell signalling network from differentiation state, but rather aid in repositioning CRC cells along the differentiation axis. Exceptions from this rule were p-S6 and p-ERK, which, though still graded along the differentiation axis, were boosted or reduced in organoid cells by individual oncogenic mutations.

Our analysis does not shed light on control of DNA replication and repair, apoptosis and absorptive lineage allocation in CRC progression, as fluctuations of markers related to these processes, namely pChk2, pH2A.X, Casp3, and KRT20 remained largely unexplained in our analysis of organoids with well-defined mutations (Fig. 4b,c). We propose that larger MC panels, and more perturbations are required to uncover how these processes shape CRC progression. Taken together, our study shows that the combination of organoid models and MC are powerful tools to dissect oncogenic signalling pathways. Several studies analysed single-cell transcriptomes of CRC organoids and tumours, showing that cell-state heterogeneity exists in the transcriptome of tumours with implications for therapy.^14,40,41^ However, the study of underlying signalling networks has only recently gained attention^9,42^ as MC was previously mainly limited to the study of extracellular markers of immune cells. Application of MC on primary cancer tissue holds the promise of dissecting signalling heterogeneity in cell-state gradients of tumour tissue. This is important as signalling pathway activities are the main targets of many modern therapies.

## Acknowledgements

The authors thank the BIH Flow & Mass Cytometry Core Facility, Gudrun Kliem (Robert Koch-Institute), and In-Fah M. Lee (Clinical Physiology at Charité) for their excellent help and technical assistance, and acknowledge funding by the German Ministry of Education and Research (BMBF), projects ZiSSTrans (02NUK047E) and MSTARS, as well as Deutsche Forschungsgemeinschaft (DFG) via the graduate programme CompCancer (RTG2424) and Sachbeihilfe MO 2783/5 (to M. Morkel).

## Data availability

All data used in this study and data analysis scripts are available on Zenodo: https://doi.org/10.5281/zenodo.6400083

## Figures

**Figure S1:**
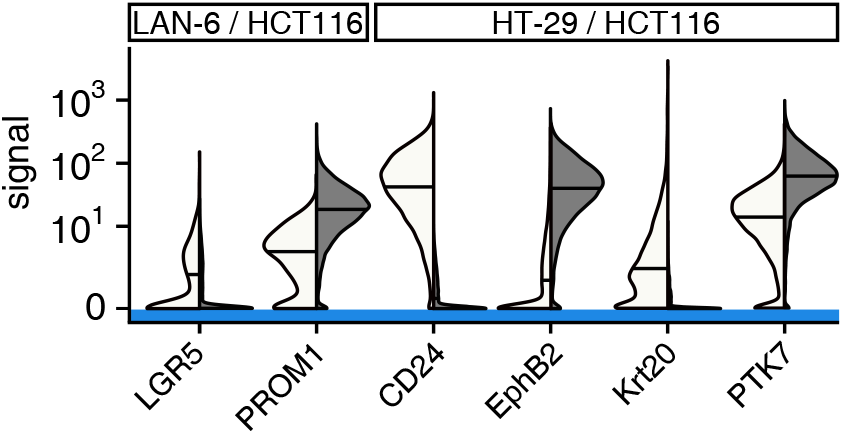
Functional control MC experiments for selected cell-state antibodies. Split violin plots with medians. Cell lines with previously known differential expressions of total proteins in question were used.

**Figure S2:**
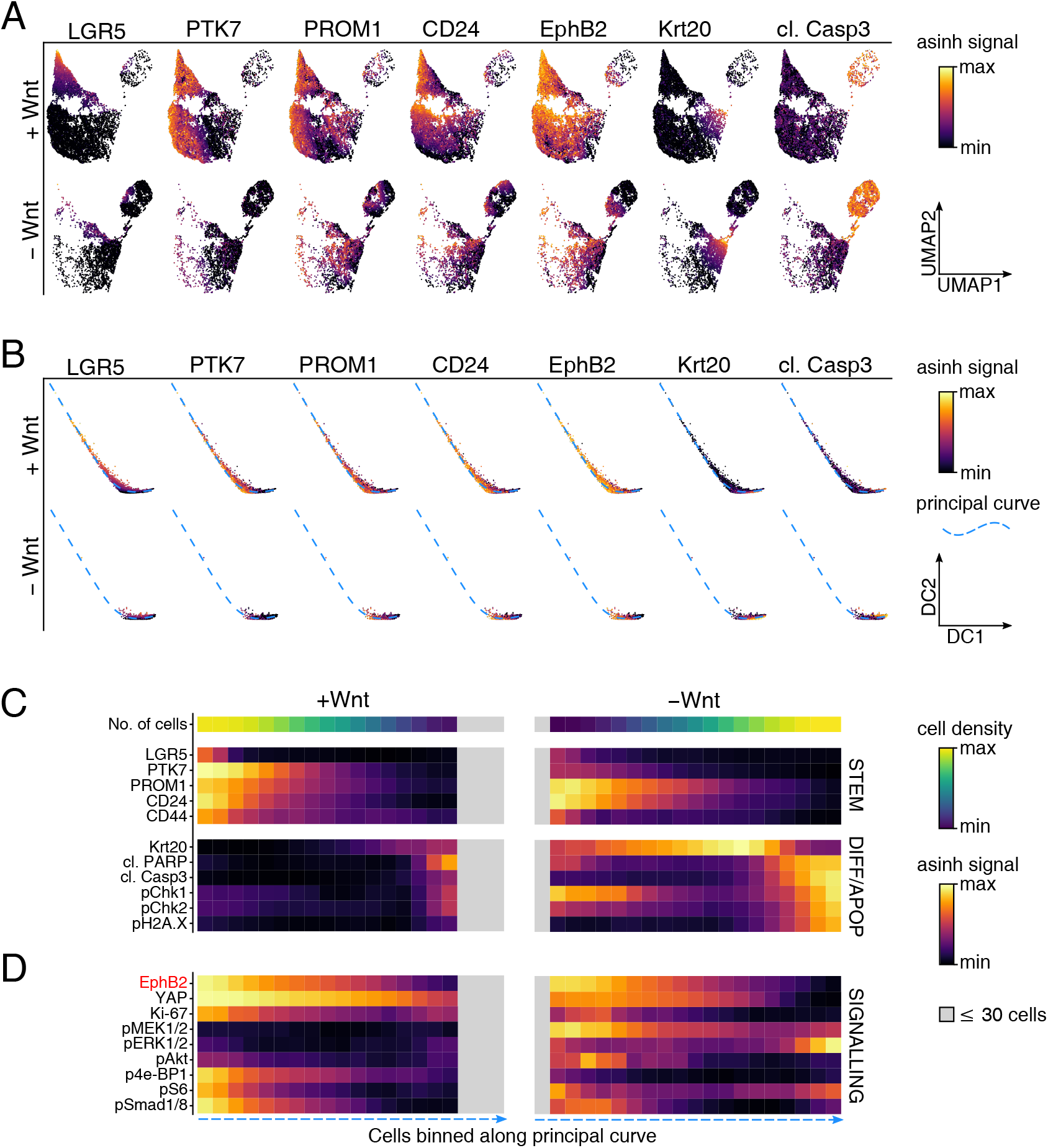
Biological replicate of Fig. 2. **a** UMAP embedding based on the 7 cell-type markers shown. Colour scale depicts normalised marker signals in ± Wnt conditions. Stem cell, differentiated, apoptotic regions are marked in red. **b** Diffusion map embedding based on the cell-type markers also used in (b). Fitted principal curve of the diffusion map space is depicted as a dashed blue line. **c/d** Heatmap showing cell distribution and mean protein marker signals within equally sized bins along the principal curve defined in (c). Means rescaled per marker and smoothed out using a 3-bin wide running average. Bins containing less than 30 cells were excluded and are shown in grey.

**Figure S3:**
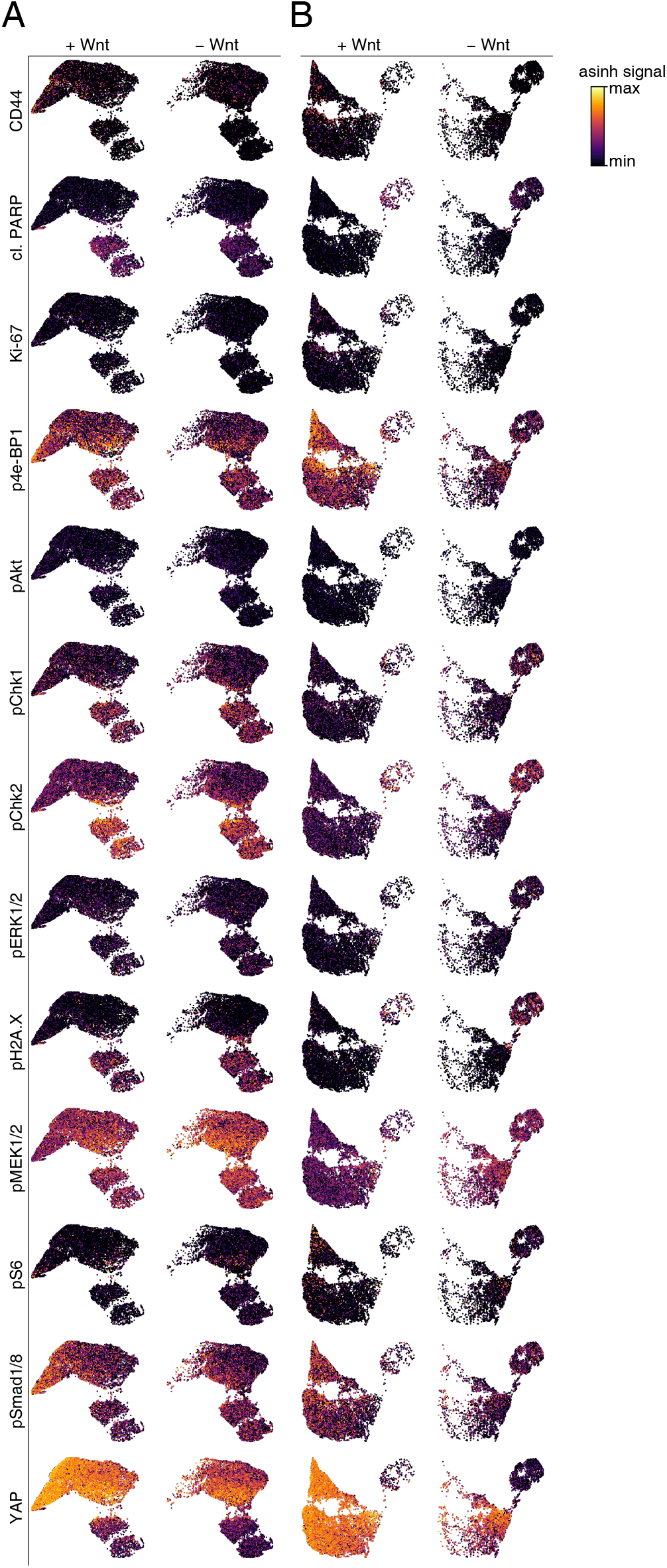
Signals of non-mapping markers for Fig. 2 & Fig. S2. **a** UMAP embedding of replicate 1 data based on 7 cell-type markers. Colour scale depicts normalised marker signals in ± Wnt conditions. **b** UMAP embedding of replicate 2 data based on 7 cell-type markers. Colour scale depicts normalised marker signals in *±* Wnt conditions.

**Figure S4:**
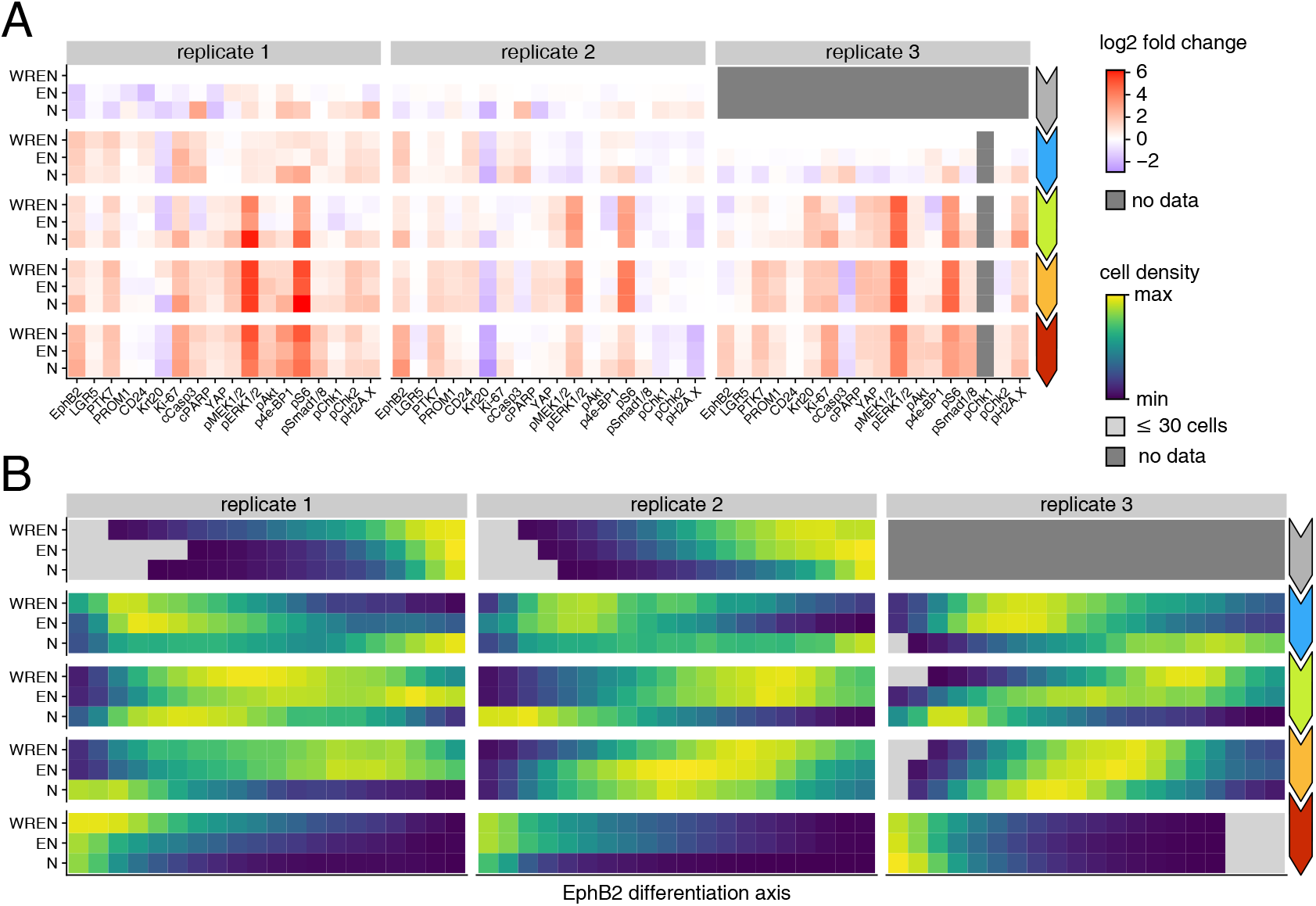
Plots of Fig. 3 split by biological replicate. **a** Log2 fold changes per marker relative to complete medium NCO (Rep. 1 & 2) or APC^ko^ (Rep. 3) condition. **b** Cell density distributions along a shared pseudo-differentiation axis defined across replicates by Ephrin B2 marker signal. Normalised by replicate, medium, & line. High Ephrin B2 (stem/TA region) on the left, low Ephrin B2 (differentiated/apoptotic region) on the right.

**Figure S5:**
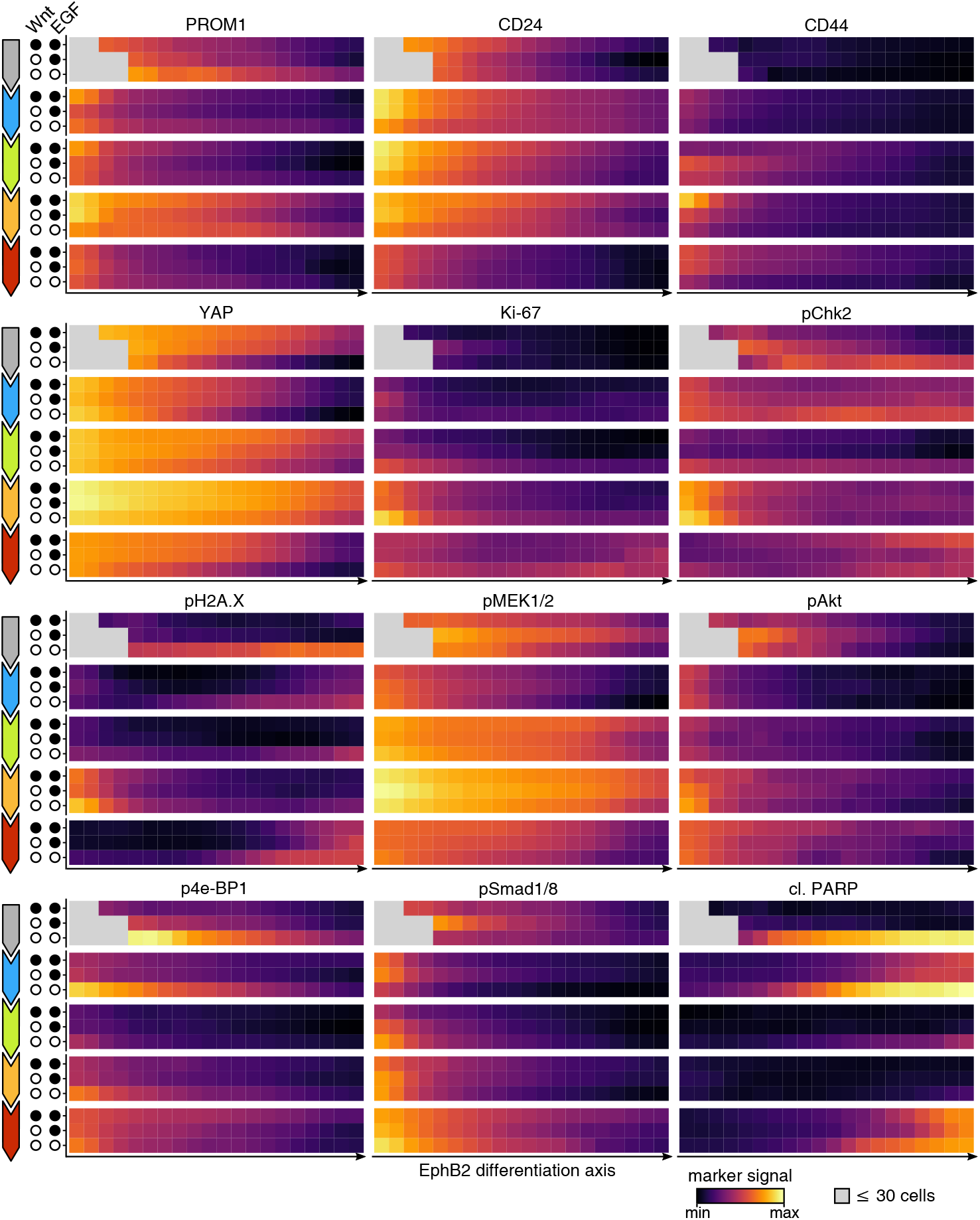
Mass cytometry signals of selected marker proteins along binned Ephrin B2 pseudo-differentiation axis defined in Fig. 3. Organoid lines as row blocks and medium conditions as intra-block rows.

**Figure S6:**
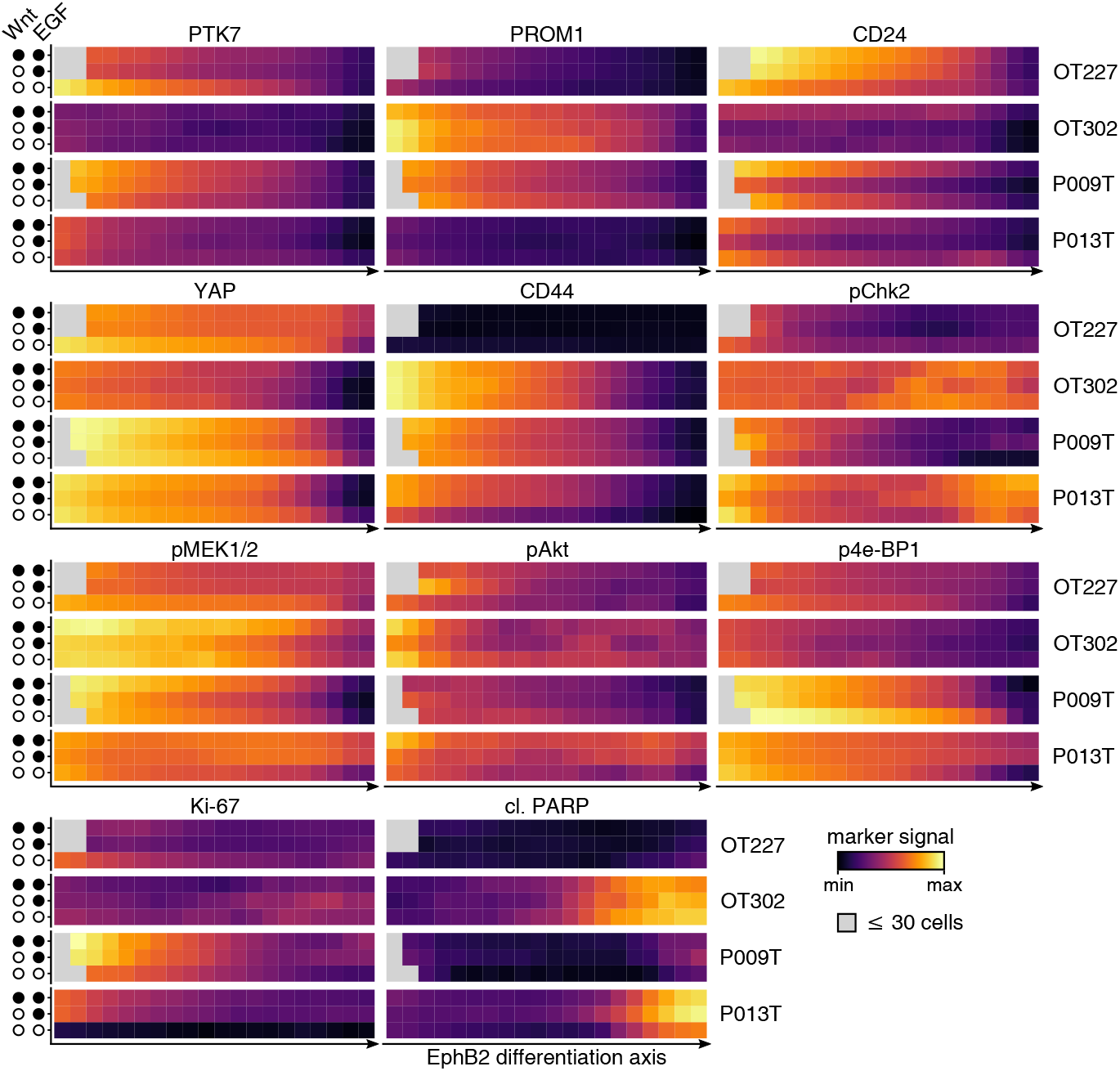
Mass cytometry signals of selected marker proteins along binned Ephrin B2 pseudo-differentiation axis defined in Fig. 5. Different patient-derived organoid lines as row blocks and medium conditions as intrablock rows.

**Figure S7:**
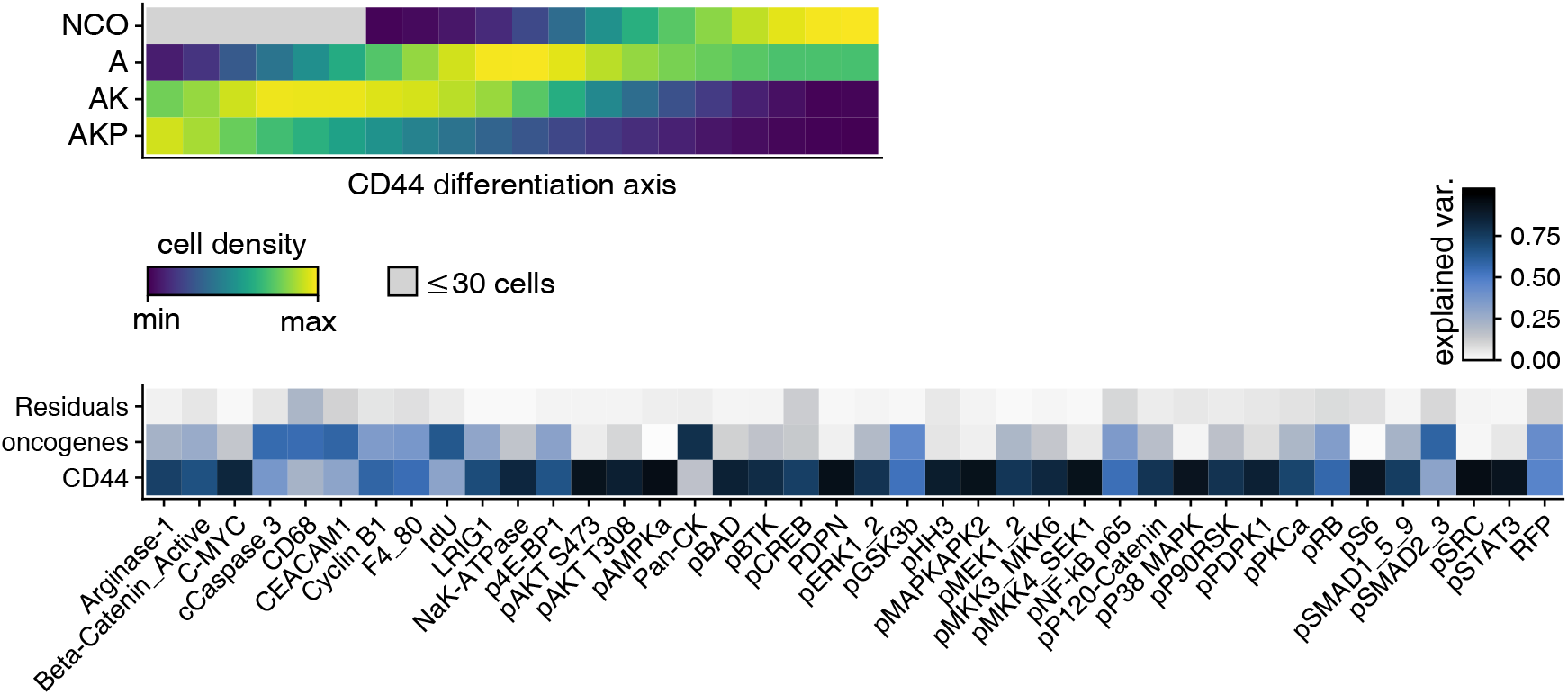
Analysis of Qin et al. 2020 mouse CRC organoid dataset. **a** Cell density distributions along a pseudodifferentiation axis defined by CD44 marker signal. Normalised by line. High CD44 (stem/TA region) on the left and low CD44 (differentiated/apoptotic region) on the right. **b** Analysis of variance per marker, using introduced oncogene and cell differentiation as described by CD44 signal as linear model factors. Colour scale shows fraction of variance explained per predictor.

